# Leaffooted bugs enrich local soil with their horizontally acquired symbiont

**DOI:** 10.1101/2025.09.03.673988

**Authors:** Bibek Singh Parajuli, John Teodosio, Alison Ravenscraft

## Abstract

Associations between hosts and their microbial symbionts are considered mutualistic when both partners benefit. While the advantages received by eukaryotic hosts from association with bacterial symbionts are frequently examined, benefits to the bacteria are rarely experimentally tested. Here, we consider whether the bug-*Caballeronia* symbiosis is truly mutualistic by measuring the effect of a leaffooted bug (*Leptoglossus phyllopus*) on the abundance of its horizontally acquired symbiont, *Caballeronia grimmiae.* We predicted that the free-living *Caballeronia* population would increase over time in the presence of its insect partner. We quantified *Caballeronia* titer in soil microcosms (i) in the presence and absence of *L. phyllopus* over time, and (ii) at different bug densities. Insect presence resulted in higher soil *Caballeronia* titer over time. As bug density increased, the soil *Caballeronia* population also increased. Additionally, soil moisture affected *Caballeronia* abundance, with moister soil supporting a larger population. These results demonstrate that the relationship between *Caballeronia* and *L. phyllopus* is truly mutualistic and add to a small but growing body of literature that has quantified the effects of eukaryotic hosts on their bacterial partners.

## Introduction

Most multicellular organisms associate with microbial symbionts (McFall-Ngai, 2014; Moran, 2007). These associations have played a crucial role in the evolution of the eukaryotic cell, evolutionary diversification, and the development of novel ecological innovations (Gerardo et al., 2020; López-García et al., 2017; McFall-Ngai, 2014; Moran, 2007; Sudakaran et al., 2017). Among the animals, insects are unique for their documented taxonomic and functional diversity of microbial symbionts (Engel & Moran, 2013; Moya et al., 2008). Benefits provided to insect hosts include nutrient synthesis, digestion of recalcitrant compounds, protection against pathogens and parasites, and increased tolerance to environmental stressors such as desiccation or extreme temperatures (Douglas, 1998, 2009; Engl et al., 2018; Flórez et al., 2015; Lemoine et al., 2020).

Insect-microbe symbioses are usually assumed to be beneficial for both partners, but most investigations focus on benefits to the host. Relatively few studies have measured *host* contributions to *symbiont* fitness (reviewed in: (Garcia & Gerardo, 2014; Hoang et al., 2024; Mushegian & Ebert, 2016; Wooldridge, 2010). Many researchers fail to consider alternative non-mutualistic possibilities- e.g., farming of symbionts as food (Mushegian & Ebert, 2016). Labeling an interaction mutualistic before the net outcome for both partners has been quantified may lead to incorrect assumptions about the interaction’s ecological and evolutionary consequences (Bronstein, 2001; Mushegian & Ebert, 2016).

Many of the benefits a microbe might receive ultimately depend on how tightly it associates with the host. Some microbes are obligately associated with insects and cannot survive separately. They are reliably transmitted from parent to offspring (vertical transmission), or between host individuals (horizontal transmission) via processes including trophallaxis (e.g. oral-oral or oral-anal feeding in social insects) (Bright & Bulgheresi, 2010; Salem et al., 2015). The advantages these obligately associated microbes receive are usually measured in terms of immediate physiological benefits, such as nutrients provided by the host (Bennett & Moran, 2015; Douglas, 1998; Fisher et al., 2017; Moran et al., 2008; Salem et al., 2015), or benefits to the host population itself, since the symbiont population is inseparable from the host population. Other microbes facultatively associate with insects and are capable of living independently (Salem et al., 2015; Wollenberg & Ruby, 2012). In addition to measuring the physiological benefits received within the host, investigators must also consider the net effect of host association on these microbes’ free-living population.

It is commonly assumed that microbes benefit from association with multicellular hosts through mechanisms including a stable, competition-free environment within the host, freedom from predation, and provisioning of nutrients to the symbiont (Garcia & Gerardo, 2014). However, these assumptions are not always correct (Garcia & Gerardo, 2014). Polyclonal symbiont populations are common in host-microbe associations, potentially leading to intense competition within the host (Baker & Romanski, 2007; Dubilier et al., 2008; Gage, 2002; Garcia et al., 2014; Martens et al., 2003). For example, in the *Wolbachia-* braconid wasp (*Asobara tabida*) symbiosis, one *Wolbachia* genotype showed decreased abundance in the presence of competing strains (Mouton et al., 2004). Moreover, while symbionts may receive protection from predation within the host, they are also subjected to the immune system (Kim, Kim, et al., 2013; Login et al., 2011; Ratzka et al., 2013). As a result, evidence suggests that in at least some symbioses, the host’s body is a comparatively stressful environment (Byeon et al., 2015; Kim, Kim, et al., 2013; Login et al., 2011; Park et al., 2018). Furthermore, although hosts can provide nutrients such as amino acids, sugars, and a carbon source to their symbiotic partner (Graf & Ruby, 1998; Ohbayashi et al., 2019; Zúñiga et al., 2018), studies showing that symbionts receiving host-derived nutrients perform better than their free-living counterparts are rare. Whether, and how often, host-provided benefits outweigh the costs of host association and lead to net positive symbiont fitness are open questions.

A few prior studies have measured how hosts affect the size of their symbiont’s free-living population. Garcia et al. (2019) tested how presence of a social amoeba, *Dictyostelium discoideum,* affected the population of two facultative symbionts, *Paraburkholderia agricolaris* and *P. hayleyella.* When the host was present, *P. agricolaris* growth was unchanged, but the abundance of *P. hayleyella* increased. In the squid-*Vibrio* symbiosis, *Vibrio* cells proliferate inside the host using host-derived nutrients and are subsequently expelled, augmenting the abundance of symbiotic *Vibrio* in the surrounding water (Graf & Ruby, 1998; Lee & Ruby, 1994). *Drosophila* fruit flies excrete N-acetyl-glucosamine and other nutrients that increase the abundance of their gut symbiont, *Lactobacillus plantarum*, in the surrounding medium (Storelli et al., 2018). Upon the death of hydrothermal vent tubeworms, their endosymbiont, *Candidatus* Endorifita, rapidly escapes the worm and reenters the environment (Klose et al., 2015). This work indicates that there are at least three mechanisms by which a host can augment its symbiont’s free-living population: via release of nutrients or via direct release of live symbiont cells into the environment, either in feces or upon host death. In all cases, increased density of the host should result in elevated local density of the free-living symbiont.

The environmentally acquired bug-*Caballeronia* symbiosis offers an opportunity to measure the net benefit (or cost) of host association for a microbial partner. Thousands of species of true bugs in at least 7 taxonomic families acquire *Caballeronia* bacteria from the soil every generation as young nymphs during their second instar (Kikuchi et al., 2007, 2011). The symbiont colonizes a specialized section of the midgut, the M4 region, which is dedicated to housing the symbiont (Itoh et al., 2019; Kikuchi et al., 2005, 2011; Ohbayashi et al., 2015; Xu et al., 2016). *Caballeronia* recycles host nitrogenous waste to produce amino acids and vitamins which the insect obtains by digesting excess symbiont cells in the anterior midgut (Ohbayashi et al., 2019; Byeon et al., 2015; Futahashi et al., 2013). This promotes faster insect development, lower mortality, and larger body size (Acevedo et al., 2021; Garcia et al., 2014; Hunter et al., 2022; Kikuchi et al., 2007; Ravenscraft et al., 2020; Stoy et al., 2023). Transcriptomic data suggest that *Caballeronia* receives sugars (ribose, rhamnose), nitrogenous wastes (allantoin, urea), and sulfonates (taurine, alkanesulphonate) as nutrients from the insect (Ohbayashi et al., 2019), and the *in vivo* titer of *Caballeronia* increases as the insect ages (Kim et al., 2014; Stillson et al., 2025). However, evidence suggests that *Caballeronia* experiences nutrient limitation in the M4, as well as exposure to antimicrobial peptides which the hosts uses to control its population, and therefore the gut is likely a stressful environment (Byeon et al., 2015; Itoh et al., 2019; Kim, Lee, et al., 2013; Lachat et al., 2024; Lee et al., 2023). Adults of some bug species can release live symbiont cells in their feces (Villa et al., 2023), but the effect on the free-living symbiont population is unknown.

We measured the effect of presence and density of the eastern leaffootted bug, *Leptoglossus phyllopus* (Coreidae), on *Caballeronia*. First, we compared soil abundances of *Caballeronia* in the presence and absence of *L. phyllopus* over time, predicting that the insect would augment the free-living *Caballeronia* population and the difference between the “insect present” and “insect absent” treatments would increase over time. In a second experiment, we quantified the abundance of *Caballeronia* in soil exposed to different host densities (0, 5, 15, and 30 bugs), predicting that increased host density would lead to higher *Caballeronia* soil titer. This study furthers our understanding of the fitness outcomes of host-microbe interactions and, thus, the conditions necessary for the persistence of these interactions.

## Methods

### *Leptoglossus phyllopus* rearing

The eastern leaffooted bug, *Leptoglossus phyllopus,* ranges from the eastern and southern United States down into central Mexico. It feeds on a wide range of seeds and fruits and can be a pest on crops including tomatoes, peppers and oranges (Mitchell, 2006). We obtained insects from a laboratory colony at the University of Texas at Arlington. The colony was established from wild individuals collected in an urban lot in Arlington in 2019. The colony was maintained in mesh cages with bush bean plants (*Phaseolus vulgaris*) and fed raw Spanish peanuts. Cages were maintained in a VIVOSUN Grow Tent (Ontario, CA) set at ∼28 °C with a 16:8 light:dark cycle. Males and females were paired to produce eggs, which were transferred to a screened plexiglass box to hatch.

### Experiment 1: Soil *Caballeronia* abundance in presence and absence of host

We constructed soil microcosms using Miracle-Gro moisture-control potting mix in 15-cm pots topped with one-gallon plastic jars with a ventilated side (Fig. 1). Peanuts attached to the side of the enclosures were provided as food. Cowpea cuttings (*Vigna unguiculata*) in floral vials were placed inside the enclosures as a source of water and shelter.

**Figure 1:**
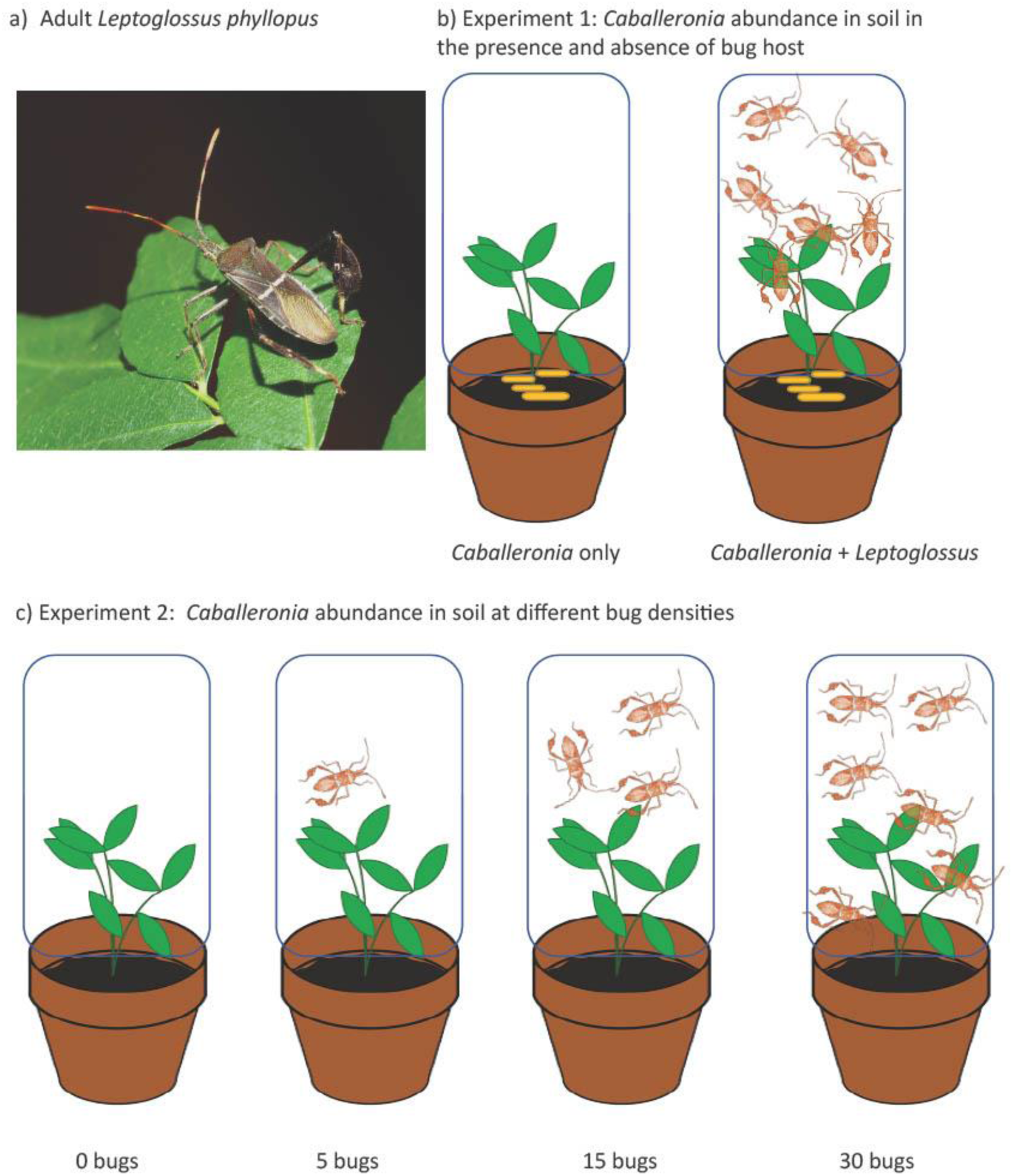
Outline of methodology. a) Adult *Leptoglossus phyllopus* b) In experiment 1, we compared soil *Caballeronia* abundance in the presence and absence of bugs. An equal number of *Caballeronia* was added to both treatments at the beginning of the experiment. In the insects-present treatment, we added 10 nymphs to each enclosure each month. We collected soil samples for 5 months. c) In experiment 2, we calculated the change in *Caballeronia* abundance over one month in soils exposed to different bug densities (0, 5, 15, and 30 bugs/microcosm).

Soil was inoculated with *Caballeronia grimmiae* isolate LepA1 (Hunter et al., 2022; Stillson et al., 2022). This strain was isolated from a laboratory colony of *Leptoglossus zonatus* at the University of Arizona, derived from insects originally collected in California, and is common in wild bugs (Ravenscraft et al., 2024). A single colony was used to inoculate 70 mL of yeast-glucose (YG) broth, which was incubated overnight at 28°C with shaking at 270 rpm. The overnight culture was diluted to an optical density (OD) of 0.8 using sterile water. For both treatments (“insect absent” and “insect present”), the soil in every enclosure was initially sprayed evenly with 2.2 ml of this cell suspension.

All enclosures were maintained in the VIVOSUN grow tent for seven months. In the *“*insect present” treatment, ten first or second-instar nymphs were placed inside each enclosure once per month. Because we observed high mortality in the second batch of added nymphs, we re-inoculated all microcosms with *Caballeronia* on Day 76. To reduce crowding, we removed eggs laid by the bugs; however, some eggs escaped our detection, resulting in accumulation of additional insects within the “insect present” enclosures.

Soil samples (∼200 mg) were collected in 1.7 ml microcentrifuge tubes at 2, 90, 108, 138, 200, and 215 days post-inoculation and held at −80 °C. Prior to sample collection, we measured soil moisture using a Dr. Meter soil moisture meter© (Hong Kong), which ranges from 1 (driest) to 10 (wettest).

### Experiment 2: Soil *Caballeronia* abundance at different bug densities

To determine whether the free-living *Caballeronia* population increases with insect density, we measured symbiont abundance in microcosms exposed to different insect densities. We constructed enclosures as described in Experiment 1. Soil in this experiment (unlike Experiment 1) was sterilized via autoclaving at 121 °C for 30 minutes and no symbiont cells were added; insects were therefore the only source of *Caballeronia*.

To colonize *L. phyllopus* nymphs with *C. grimmiae*, Lep1A1 cells were cultured at 28 °C in YG broth overnight, shaking at 270 rpm. To obtain log-phase cells, 500 μL of the overnight culture was transferred to 2.5 mL of fresh broth and cultured for several hours. Log phase culture was diluted to an OD of 0.1 and fed to freshly molted second instar nymphs for three days. Nymphs were then introduced into enclosures at different densities (0, 5 bugs, 15 bugs, and 30 bugs). Enclosures were maintained in the VIVOSUN tent for 33 days. Soil moisture and bug density were monitored weekly. All nymphs had either developed into adults or died by the end of the experiment. Soil samples were collected from each enclosure at the beginning and end to quantify *Caballeronia*.

Additionally, at the end of the experiment one soil sample per enclosure was treated with PMAxx™ (Biotium, California, US) to quantify viable *Caballeronia.* PMAxx binds to DNA and prevents its amplification, but cannot penetrate intact cell membranes; it thereby prevents amplification of DNA from dead or dying cells. We used the manufacturer’s protocol with slight modifications: We treated each soil with a 100 nM PMAxx instead of [manufacturer’s recommended concentration], followed by incubation in the dark for 15 minutes. Soil suspensions were exposed to blue light for 30 minutes with intermittent shaking to deactivate the PMAxx.

### DNA extraction and qPCR

DNA was extracted from the soil samples using Quick-DNA™ Fecal/Soil Microbe Miniprep kit (Zymo Research) following the manufacturer’s protocol for soil, except that we used a TissueLyser II at 25.0 Hz for 7 minutes to homogenize the samples instead of the proprietary bead beater. Additionally, to improve DNA yield, we added 800 μL of Genome lysis buffer and 400 μL of 100% ethanol to the filtrate from the Zymo-Spin III-F filter.

To quantify symbiont titer, we designed species-specific primers using the genome of *C. grimmiae* Lep1A1. We identified a single copy gene unique to this species (Stillson et al., 2022) and used PrimerQuest™ Tool (IDT) to design primers qdhm-F (5’-TTGCGACCTGTTCCTTTCA-3’) and qdhm-R (5’-CGTGTGATAGTCGCCGTTAT-3’), which target a 151-bp region of the dihydromethanopterin reductase (DHMR) gene. NCBI BLAST predicted *C. grimmiae* Lep1A1 to be the sole organism amplified by these primers, and we experimentally verified primer specificity by running diagnostic PCR with several phylogenetically distinct strains of *Caballeronia* (LP003, LZ003, LZ029) and a closely related *Cupriavidus* strain (LZ004). We tested a gradient of annealing temperatures from 60-68 °C and the temperature that produced the brightest band (60 °C) was used.

To create standards for absolute quantification of DHMR copy number, we amplified Lep1A1 genomic DNA with the qdhmR/F primers. The concentration of the amplified PCR product was determined using a Qubit dsDNA High-Sensitivity Assay (ThermoFisher Scientific, Waltham, MA, USA). We calculated the number of DHMR copies using the DNA Copy Number and Dilution Calculator (ThermoFisher Scientific). We made 10-fold dilutions ranging from 10 copies per μL to 10^7^ copies per uL.

The qPCR assays were run on an Applied Biosystems 7300 Real-Time PCR System (ThermoFisher Scientific). Each reaction contained 10 μL PowerUp SYBR Green master mix (Applied Biosystems, Waltham, Massachusetts), 4 μL of molecular-grade water, 2 μL each of forward (500 nM) and reverse primers (500 nM), and 2 μL of template DNA. The thermocycling program was 50 °C for 2 minutes to activate uracil-DNA glycosylase (which prevents reamplification of carryover PCR products), 95 °C for 2 minutes, followed by 40 cycles of 95 °C for 15 seconds and 60 °C for 1 minute. To determine if non-specific products were amplified, we finished with a melt curve ramping from 60°C to 95°C. Each soil sample was run in triplicate and replicates were averaged. No template controls were included on every run. Amplification efficiency ranged from 90% to 116% across all runs.

### Analysis

In our first experiment, we analyzed the effect of *L. phyllopus* presence on the abundance of *Caballeronia* in soil microcosms over time with linear mixed effect models (LMM) using the lmer command in R. A full model was run with fixed terms for treatment (insects present or insects absent), number of days post soil inoculation (time), scaled squared days post inoculation (time^2^) to account for a nonlinear relationship, and soil moisture, as well as all pairwise interactions between these terms except for the time x time^2^ interaction. Microcosm was accounted for as a random effect. We performed backward model selection with likelihood ratio tests to determine the final model. Model residuals were tested for normality and homoscedasticity using the Shapiro-Wilk test and by plotting the residuals against the predicted values, respectively. *Caballeronia* abundance was log-transformed to meet the normality assumption of linear mixed-effect models.

For the second experiment, we used a linear regression model (lm command in R) to determine the effect of bug density on the *Caballeronia* population after a one-month exposure to insects. Independent variables were bug density and bug density squared (to allow for a non-linear relationship). The dependent variable was the change in *Caballeronia* DHMR copy number from the initial time point (0 days) to the final time point (33 days). Prior to calculating the difference, we applied a ln(mean + 1) transformation to DHMR copy number. To assess whether DNA came from living cells, we also tested for correlation between DHMR copy numbers in replicated soil samples from each enclosure at the final timepoint that were, or were not, treated with PMAxx.

## Results

### Experiment 1: *C. grimmae* abundance in soil over time is higher in the presence of *L. phyllopus*

We inoculated soil microcosms with cultured *Caballeronia* cells and compared the soil *Caballeronia* population with or without *L. phyllopus*. We predicted that insects would supplement the free-living symbiont population such that over time, soil exposed to insects would contain more and more *Caballeronia* compared to unexposed soil. Unexpectedly, symbiont populations decreased in both treatments over the course of the 215-day experiment, likely because our inoculations overshot the soil’s carrying capacity for *Caballeronia* (Fig. 2). However, this decrease trended slower in soils exposed to bugs, resulting in higher *Caballeronia* populations in the microcosms with insects (treatment*time interaction term: df=1, χ²=2.8, p-value=0.09). Although the interaction between insect presence and time was only marginally significant, this was likely due to comparatively small sample size (15 microcosms per treatment) and high variance: We could not track where insects defecated or died, but these factors likely caused fine-scale but high-magnitude variation in *Caballeronia* density in the soil. We retained the interaction in the final model because omitting it resulted in flawed predictions. Specifically, the model with only main effects of treatment and time predicted that *Caballeronia* population would always be higher in soils exposed to insects, even at the beginning of the experiment, prior to insect introduction. However, all soils were inoculated identically, and we confirmed that *Caballeronia* populations at the start of the experiment were equal between microcosms with and without bugs (t-test, t (27.43) = −0.1486, p=0.883). Furthermore, *Caballeronia* abundance at day 90, after we re-inoculated all microcosms, was also equivalent across treatments (t-test, t(25.509) = 0.50452, p = 0.62). Therefore, the increased abundance of *Caballeronia* in the presence of insects was driven by later time points.

**Figure 2:**
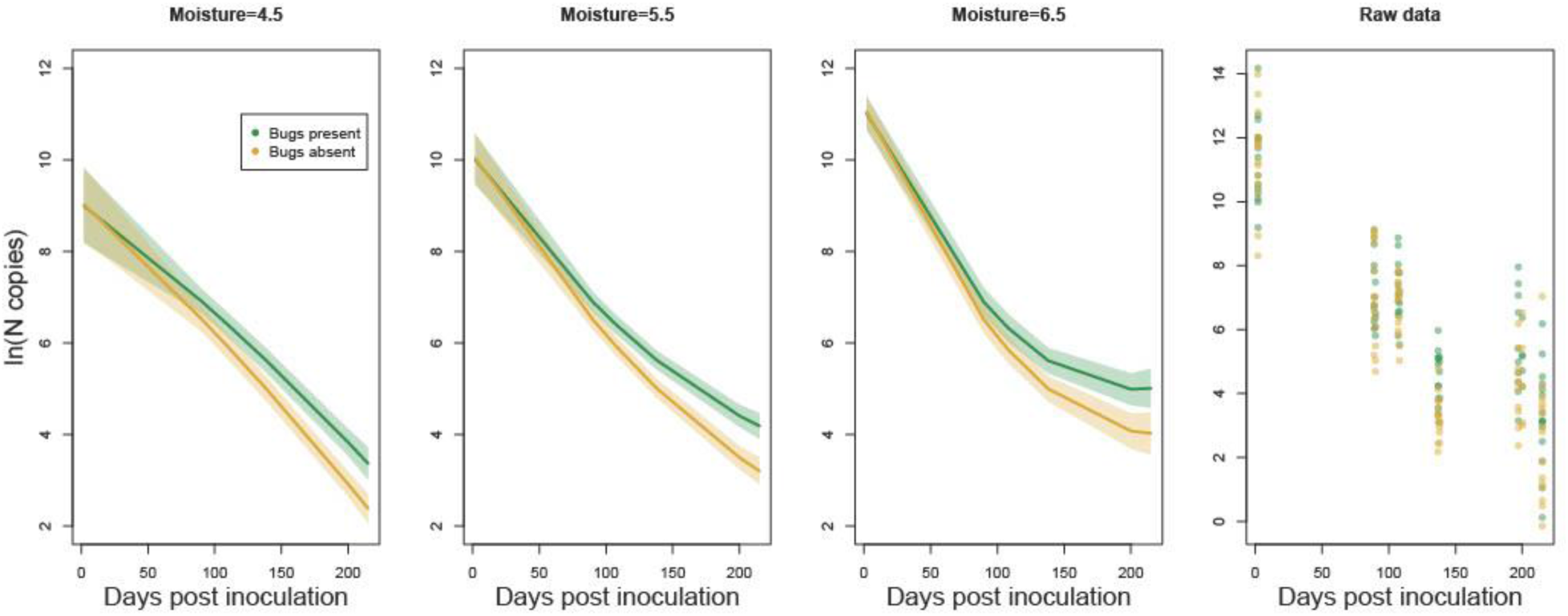
(A-C) Predicted effect of *Leptoglossus phyllopus* on *Caballeronia grimmiae* abundance in soil over time at different moisture levels. Colored lines show model estimated means and shading depicts estimated standard errors. A) Moisture level=4.5 B) Moisture level=5.5 C) Moisture level= 6.5 D) Raw data points for *Caballeronia* abundance in soil over time. Each point reports the mean of triplicate titer measurements. Y-axis reports the titer of *C. grimmiae* in 200 mg of soil.

Interestingly, we also found that wetter soils supported larger *Caballeronia* populations. This relationship was nonlinear and depended on time, with *Caballeronia* population predicted to decrease to extinction in drier soil, but to stabilize in wetter soil (Fig. 2; moisture*time interaction: df=1, χ²=, p=0.002; moisture*time^2^ interaction: df=1, χ²=, p=0.002). Soil moisture did not systematically differ between microcosms with and without bugs (model predicting moisture as a function of time and treatment, with random effect for microcosm: time*moisture interaction, df= 1, x=0.76, p=0.4).

At 125 days post-inoculation and with average soil moisture of 5.5, the model estimated there were about 1,000 more *Caballeronia* copies per gram of soil in microcosms with bugs than without bugs.

### Experiment 2: *Caballeronia* abundance in soil increases with bug density

Next, we tested whether higher insect density promotes higher *Caballeronia* abundance in soil. For this experiment, we used autoclaved soil and only the insects (not the soil) were inoculated with *Caballeronia*. Over the course of one month, *Caballeronia* titer remained essentially unchanged in insect-free microcosms, but increased with rising insect density (Fig. 3; bug density: df=1, χ²=58, p<0.001; bug density^2^: df=1, χ²=34, p=0.004). The benefit to *Caballeronia* showed diminishing marginal returns: At and above 15 bugs per microcosm, the one-month increase in *Caballeronia* abundance stabilized at about 280 additional copies per gram of soil.

**Figure 3:**
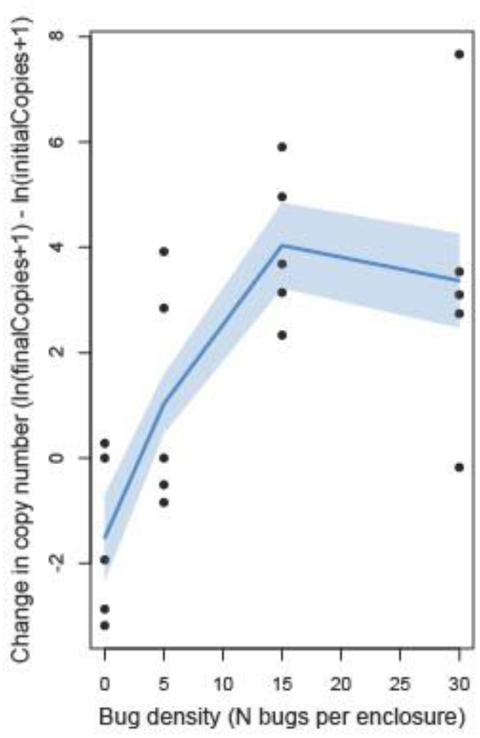
Effect of host density on change in *Caballeronia grimmiae* abundance in soil over one month. The blue line shows the model estimated mean difference and blue shading depicts the estimated standard error. Each data point is an observed value (difference between the means of triplicate titer measurements).

To ascertain whether *Caballeronia* were viable, we collected duplicate soil samples from each microcosm at the end of the experiment and treated these with PMAxx, which prevents amplification of DNA from dead cells. Estimates of live *Caballeronia* density (PMAxx-treated soils) did not differ from estimates of total *Caballeronia* density in the same soils (without PMAxx treatment) (Fig. S1; LRT: df=1, χ² =2.4552, p > 0.05), suggesting that essentially all the *Caballeronia* we detected were viable. However, we note that PMAxx is not designed for use with soil samples and therefore, our estimates of live cell count may be imperfect.

After one month, our model predicted a decrease of four *Caballeronia* cells per gram of soil in the absence of bugs, versus an increase of 280 cells per gram after exposure to 15 or more bugs (Fig. 3).

## Discussion

We found that both the presence and population density of leaffooted bugs were correlated with increased abundance of their bacterial symbiont in the environment. In our first experiment, pre-inoculated soil exposed to insects over a period of seven months contained more *Caballeronia* than soil unexposed to insects. In a second experiment using uninoculated soil, *Caballeronia* abundance increased with insect density; the magnitude of increase started to plateau around 15 bugs and *Caballeronia* received diminishing marginal returns from additional insects.

These findings suggest that the free-living *Caballeronia* population does benefit from association with *Leptoglossus*. This could result from three mechanisms. First, bugs may excrete live cells in frass. Squash bug (*Anasa tristis)*, another leaf-footed bug (Coreidae), releases live *Caballeronia* in its feces (Villa et al., 2023). While we haven’t quantified the release of *Caballeronia* from *L. phyllopus*, we have previously isolated live symbiont from the insects’ feces, and live cells have been detected in the feces of the congener, *L. zonatus* (L. Sullivan, personal communication, February 2024). Second, cells may escape from dead insect carcasses. In dead mammals, collapse of the immune system leads to rapid proliferation of the gut microbial population (Heimesaat et al., 2012), some of which may be released into the environment. Similar proliferation might occur in dead bugs, since production of the antimicrobial peptides used to control *Caballeronia* in the M4 (Byeon et al., 2015; Futahashi et al., 2013) should cease. (However, *Caballeronia* are aerobic and oxygen could rapidly deplete in a decomposing insect.)

In both of our experiments, we left dead insects in the microcosm to decompose. Enrichment of the symbiont population via escape from dead hosts has been observed in tube worms: A medium-sized worm releases up to 7 × 10^5^ *Candidatus* Endoriftia symbionts into the environment upon its death (Klose et al., 2015). Third, leaffooted bugs might release nutrients into the soil from frass or dead insect bodies, which can increase soil microbial biomass (Yang & Gratton, 2014). The deposition of dead cicada carcasses in forest plots increases soil nitrate and ammonium availability, leading to a higher relative abundance of soil bacteria (Yang, 2004). *Drosophila* larvae promote persistence of their facultative symbiont *Lactobacillus plantarum* in their shared habitat through intestinal secretions of the nutrient N-acetyl-glucosamine (NAG) (Storelli et al., 2018). Whether cells or nutrients released from true bugs can similarly affect the population of free-living *Caballeronia* should be further investigated.

In addition to insect presence and density, soil physical and chemical properties can impact environmental bacterial abundances. Various studies have demonstrated that wetter soil conditions are correlated with an increased abundance and activity of soil microbial communities (Colombo et al., 2016; Manzoni et al., 2012; Morugán-Coronado et al., 2019; Zhang et al., 2013). Similarly, we found that moister soil was associated with higher *Caballeronia* abundance. Furthermore, the observed decline in symbiont population in dry soil suggests that insects may act as a refuge for *Caballeronia* in drier conditions. Further investigation on how soil moisture and chemistry (e.g. pH) affect host acquisition of *Caballeronia* will prove valuable in understanding how environmental conditions affect insect-microbe interactions.

Interestingly, in experiment 1 *Caballeronia* abundance declined over time in both insect-present and insect-absent treatments (Figure 2), most likely due to our initial inoculation exceeding the soil’s carrying capacity. In dry soils, *Caballeronia* population trended towards extinction, but in wetter soils, the population appeared to stabilize at a carrying capacity of approximately 5.7 x 10^3^ cells per gram without insects and 1.0 x 10^4^ cells per gram with insects (Fig. 2c). In contrast, a previous study estimated the natural abundance of *Caballeronia* in agricultural fields to be 1.1 x 10^7^ copies per gram of soil (Tago et al., 2014). This difference is likely due to different soil physical and chemical conditions, as discussed above. Most notably, we used commercial potting soil while Tago and colleagues quantified *Caballeronia* in agricultural soil. Regardless, we still observed a positive effect of insect presence on the free-living population, with the population exposed to bugs decreasing more slowly over time.

Only a handful of studies have measured how hosts affect the size of their symbionts’ free-living population. In the legume-rhizobium symbiosis, nitrogen-fixing bacteria are acquired from the soil and reproduce within root nodules, eventually escaping, repopulating the soil, and infecting future generations (Denison & Kiers, 2011). Indeed, one study found the soil rhizobial population to be five times larger in plots where leguminous plants were present (Kuykendall, 1989). In ocean environments, association with bobtail squid allows the bioluminescent bacterium *Vibrio fisheri* to grow to higher population sizes due to competition-free growth within the host’s symbiotic organ, followed by daily release of bacteria into the environment (Lee & Ruby, 1994; Wollenberg & Ruby, 2012). Enrichment of the local environment with symbionts should increase the likelihood that offspring find a symbiotic partner, while leaving ample opportunity for offspring to acquire other symbiont strains. This allows for the possibility that offspring acquire locally adapted strains that confer enhanced fitness under local conditions, while still bet-hedging against the risk that offspring acquire an inferior symbiont, or fail to acquire any symbiont (Bruijning et al., 2021; Lange et al., 2023; Stillson et al., 2025).

While we were primarily interested in the effect of the host on symbiont fitness, supplementing the free-living symbiont is also likely to benefit the host. In the first experiment, the final difference of ∼4,300 cells per gram represented a ∼75% increase compared to soils unexposed to insects. In the second experiment, *Caballeronia* was estimated to decrease by 4 cells per gram of soil over the course of one month in the absence of bugs but increased by about 280 cells per gram in the presence of at least 15 bugs (Fig. 3). These increases are likely meaningful for insect fitness, since in another *Caballeronia*-hosting bug, *Riptorus pedestris*, it took just 80 *Caballeronia* cells to infect an average of 50% of nymphs and 3,500 cells to infect 100% of nymphs (Kikuchi & Yumoto, 2013).

We studied the effect of the host on one symbiont strain, but host effects might vary across different host-symbiont pairings. For example, the advantage conferred by host association to microbial symbionts varies among symbiont species in the social amoeba (*Dictyostelium discoideum*)-*Paraburkholderia* symbiosis*. Parabukholderia hayleyella* increased in population when associated with *Dictyostelium,* whereas *P. agricolaris* decreased when the host was present (Garcia et al., 2019). Different insect species might also have different effects on environmental symbiont abundance: While it seems likely that many coreid insects augment local populations of *Caballeronia* via release of cells in frass, the alydid bean bug (*Riptortus pedestris*) never sheds live *Caballeronia* in its frass. These observations suggest that outcomes of the relationship between true bugs and the *Caballeronia* can vary with different combinations of host and symbiont strains. Additionally, the magnitude and direction (e.g. beneficial or detrimental) of these outcomes for one or both partners can shift based on biotic, abiotic, and stochastic contexts (e.g. priority effects) (Chen, Junker, et al., 2024; Chen, Kwong, et al., 2024; Leung & Poulin, 2008; Moxon & Kussell, 2017; Stillson et al., 2025).

## Conclusion

While the impact of microbial symbionts on host fitness has been the focus of most symbiosis research (Cani, 2018; Douglas, 1998; Kaltenpoth & Flórez, 2020; McFall-Ngai, 2002), benefits conferred to the symbiont are rarely quantified (Garcia & Gerardo, 2014; Hoang et al., 2024). We found that the population of free-living *Caballeronia* increases in the presence of its insect host, *Leptoglossus phyllopus*. Thousands of true bug species depend on *Caballeronia* for their nutrition; our findings suggest that these relationships can benefit the bacteria as well as the insects. Our results are consistent with similar studies of other environmentally acquired symbioses, including the squid-*Vibrio* luminescent symbiosis and the social amoeba-*Paraburkholderia* nutritional symbiosis (Garcia et al., 2019; Lee & Ruby, 1994). It is important to understand the effect of the multicellular hosts on their microbial partners to better understand the evolutionary and ecological dynamics underlying host-symbiont relationships.

## Data Availability

Data and the R analysis script are available on figshare.

https://figshare.com/articles/dataset/Effect_of_Leptoglossus_phyllopus_on_Caballeronia_abundance/30034696

## Supplemental figures

**Supplemental figure 1:**
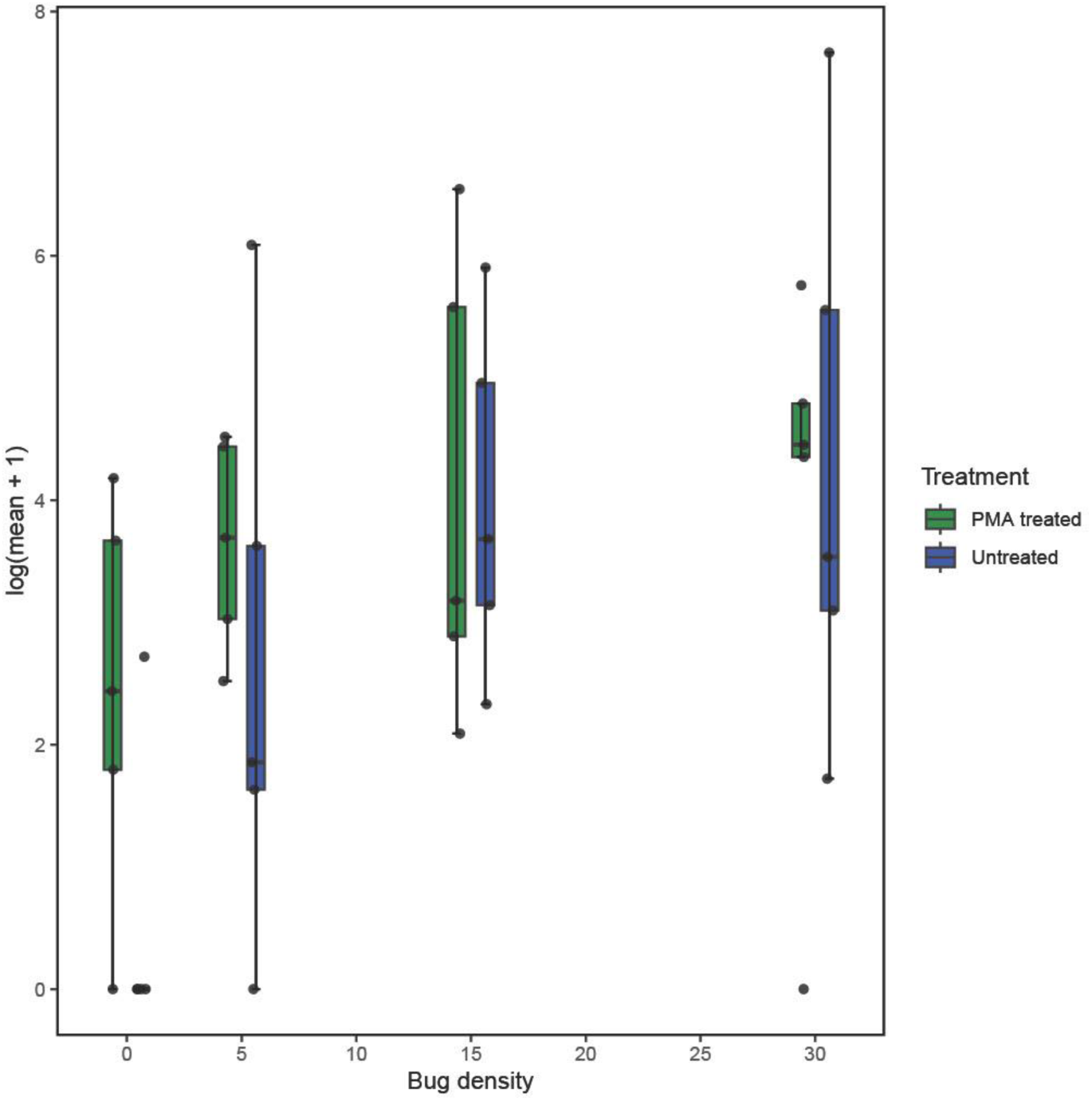
Comparison between live and total *Caballeronia* titers in soil across different bug densities at the end of Experiment 2. Two soil samples were taken per microcosm and one from each set was treated with PMAxx to prevent quantification of DNA from dead *Caballeronia* cells. There was no significant difference in titers between the PMAxx treated and untreated soil samples.

